# Alterations in brain network organization in adults with obesity as compared to healthy-weight individuals and seniors

**DOI:** 10.1101/685081

**Authors:** J. Ottino-González, H.C. Baggio, M.A. Jurado, B. Segura, X. Caldú, X. Prats-Soteras, C. Tor, M.J. Sender-Palacios, N. Miró, C. Sánchez-Garre, M. Dadar, A. Dagher, I. García-García, M Garolera

## Abstract

Life expectancy and obesity rates have drastically increased in recent years. An unhealthy weight is related to long-lasting biological deregulations that might compromise the normal course of aging. The aim of the current study was to test whether the network composition of young adults with obesity would show signs of premature aging. To this end, subjects with obesity (N = 30, mean age 32.8 ± 5.68), healthy-weight controls (N = 33, mean age 30.9 ± 6.24) as well as non-demented seniors (N = 30, mean age 67.1 ± 6.65) all underwent a resting-state MRI acquisition. Functional connectivity was studied by means of graph-theory measurements (i.e., small-world index, clustering coefficient, characteristic path length, and mean degree). Contrary to what expected, obesity in adults was related to disruptions in small-world properties driven by increases in network segregation (i.e., clustering coefficient) as compared to elders. Also, this group showed alterations in global and regional centrality metrics (i.e., degree) relative to controls and seniors. Despite not mimicking what was here shown by seniors, the topological organization linked to an obesity status may represent a flaw for cognitive functions depending on the rapid combination between different modular communities.

## 1. INTRODUCTION

Our life expectancy is longer than it has ever been before. Above 30% of the population from Western developed countries will be older than 60 years by 2050 (World Health Organization, 2015). For this reason, research on the psychobiological factors that have an impact over the so-called “successful aging” is rapidly gaining popularity. One of such factors is obesity. The World Health Organization (WHO) estimates that the prevalence of obesity has tripled in the last decades (WHO, 2017). The far-reaching consequences of this medical condition involve several chronic diseases, such as type II diabetes, cardiovascular disorders and cancer (Wade, Carslake, Sattar, Davey Smith, & Timpson, 2018). On a similar note, obesity could accelerate brain aging (Ronan et al., 2016; Tzanetakou, Katsilambros, Benetos, Mikhailidis, & Perrea, 2012), increasing the odds of suffering late-onset dementia (Bischof & Park, 2015). Neuroanatomical studies suggest that obesity is associated with changes in gray and white matter tissues (García-García et al., 2018; Repple et al., 2018; Zhang et al., 2018) that could resemble those prompted by aging.

The functional organization of the brain is intricate. Distinct and anatomically distant regions interact with each other overall contributing to cognitive activity. Graph-based indexes could be useful in disentangling the complex rendering of large-scale networks. Notably, one of the most explored metrics is the small-world (SW) index. The SW index reflects the inherent architecture of real-world scenarios to which the brain is not oblivious to. This type of network organization is especially suited for cognitive function as it facilitates the presence of highly-specialized modules exchanging information at a minimal cost (Rubinov & Sporns, 2010). This is computed as the ratio of segregation and integration measures. The clustering coefficient, a surrogate of segregation, represents how neighboring nodes tend to aggregate together to presumably process specific sources of information altogether (Rubinov & Sporns, 2010). The characteristic path length, or the shortest path between modules, offers instead a measure of integration. Hence, the fewer the jumps, the better the communication between remote communities is (Rubinov & Sporns, 2010). The SW index consequently reflects the proportion of short (i.e., intra-modular) and long-range connections (i.e., inter-modular). A network is considered as SW when returns a fraction slightly favoring the former (i.e., segregation > integration). Any deviation from this ratio could suggest a shift towards a more random or more regular type of network. Typically, normal and pathological aging are linked to alterations in SW properties primarily driven by drops in segregation metrics (Geerligs, Renken, Saliasi, Maurits, & Lorist, 2015). Other graph-related parameters often exploited are those reflecting centrality. Particularly, the mean degree, echoes the number of effective connections within the network (Rubinov & Sporns, 2010). The aging process is mainly correlated with a progressive trimming of intra-modular connections (Geerligs et al., 2015; Sala-Llonch, Bartrés-Faz, & Junqué, 2015; Sala-Llonch et al., 2014) which illustrates loses in network segregation.

To date, and to the best of our knowledge, only four studies have explored the brain functional organization in obese individuals with graph-theoretical parameters. The first two reported lower segregation in individuals with obesity when compared to healthy-weight subjects (Baek, Morris, Kundu, & Voon, 2017; Chao et al., 2018). The third and fourth study described global brain connectivity (i.e., number of effective connections) decreases primarily among prefrontal circuits (Geha, Cecchi, Todd Constable, Abdallah, & Small, 2017; Moreno-Lopez, Contreras-Rodriguez, Soriano-Mas, Stamatakis, & Verdejo-Garcia, 2016). As mentioned above, disturbances within SW properties as well as alterations in centrality degree are common signatures of aging. Thus, their presence in young adults with obesity might be symptomatic of premature aging. Because of its potential long-term consequences over health, the aim of the current study was to compare the network topology of adults with obesity to otherwise healthy elders by means of graph-theoretical measures. Our hypothesis is that an obesity status could precipitate the aging of the brain. Whereas we further expect analogous network profiles between adults with obesity and seniors, namely droppings in measures of network segregation and centrality as compared to healthy controls.

## 2. METHODS

### 2.1. Participants

Two independently collected samples were included in this work. One sample comprised 30 stroke-free non-demented seniors (56-85 years old). The recruitment procedure of this group is fully described in Abós et al. (2017). Briefly, exclusion criteria consisted on presence of psychiatric or neurological comorbidity, low global intelligence quotient (IQ) estimated by the WAIS-III Vocabulary subtest score (> 7 scalar score) (Wechsler, 1999), and a Mini-Mental state examination score below 25 (Folstein, Folstein, & McHugh, 1975). For the current work, senior participants with obesity (body mass index [BMI] equal to or higher than 30 kg/m2) were excluded (N = 4) as well as those presenting MRI pathological findings unrelated to the course of normal aging (i.e., white matter hyperintensities, N = 2). Eligible subjects underwent a magnetic resonance imaging (MRI) acquisition at the *Hospital Clínic de Barcelona*.

The other sample included 66 adults (21-40 years old) from public health care centers belonging to the *Consorci Sanitari de Terrassa*. This sample was recruited in the context of a broader line of research revolving around obesity and brain function. Part of this sample was used in previous works (Ariza et al., 2012; Caldú et al., 2019; García-García et al., 2013a, 2013b, 2013c, 2015; Marqués-Iturria et al., 2013, 2014, 2015; Ottino-González et al., 2017, 2018, 2019). In accordance to the criteria established by the WHO (2012), thirty adults were considered as obese and 33 as normal weight (BMI < 24.9 kg/m^2^). We excluded participants with any cardiometabolic neurological or psychiatric comorbidities. Individuals underwent three visits. The first consisted of a fasting blood sample extraction and a medical exploration. The second visit included an extensive neuropsychological evaluation ruling out subjects with a Vocabulary subtest scalar score below 7 (WAIS-III or WISC-IV) (Wechsler, 1999, 2007). The third and last visit comprised an MRI acquisition at the *Hospital Clínic de Barcelona*.

Additionally, participants from all three groups exhibiting excessive head motion were discarded (N = 16). Excessive head movement was defined as (1) mean interframe head motion greater than 0.3 mm translation or 0.3° rotation, and (2) maximum interframe head motion greater than 1 mm translation or 1° rotation.

### 2.2. MRI acquisition

Images were acquired with a 3T Siemens scanner (MAGNETOM Trio, Siemens, Germany). Each participant underwent a T1-weighted structural image for co-registration and parcellation purposes with the MPRAGE-3D protocol [echo time (TE) = 2.98 ms; repetition time (TR) = 2300 ms; inversion time = 900 ms; 256 mm field of view [FOV]; matrix size = 256 × 256; voxel size = 1.0 × 1.0 × 1.0 mm^3^). Resting-state volumes were collected using a multi-slice gradient-echo EPI sequence covering the whole brain (TE = 19 ms; TR = 2000 ms; 3 mm slice thickness; 90° flip angle; 220 mm FOV; voxel size = 1.7 × 1.7 × 3.0 mm^3^).

### 2.3. Image pre-processing

Basic functional image preprocessing, using AFNI (http://afni.nimh.nih.gov/afni) and FSL (https://www.fmrib.ox.ac.uk/fsl) tools, included: discarding the first five volumes to allow magnetization stabilization, despiking, motion correction, brain extraction, grand-mean scaling (to keep signal variation homogeneous across subjects), linear detrending, and high-pass filtering (> 0.01 Hz). Functional images and T1-weighted volumes were co-registered and then non-linearly normalized to the MNI ICBM152 template. Then, images were non-linearly transformed to MNI space at 3 × 3 × 3 mm^3^ voxel size using SPM (http://www.fil.ion.ucl.ac.uk/spm/).

To remove the effects of head motion and other non-neural sources of signal variation from the functional data sets, we used an independent component analysis (ICA)-based strategy for Automatic Removal of Motion Artifacts (ICA-AROMA) (Pruim et al., 2015). This method uses individual resting-state data to perform ICAs and automatically identify artifact-related independent components. The time courses of components considered as artifactual were stored to be used as regressors during network computation.

The six motion parameters obtained from the realignment procedure, as well as the average white matter and cerebrospinal fluid signals were kept as regressors in network reconstruction. To generate the corresponding white matter and cerebrospinal fluid (lateral ventricle) masks, T1-weighted structural images were segmented using FreeSurfer (https://surfer.nmr.mgh.harvard.edu/). The resulting binary masks were linearly transformed from structural to native functional space using FSL-FLIRT (https://fsl.fmrib.ox.ac.uk/fsl/fslwiki/FLIRT). To prevent these masks from extending to the adjacent gray matter due to resampling blur in linear transformation, white matter masks were then “eroded” by applying a threshold of 0.9, while ventricular masks were thresholded at 0.3.

### 2.4. Connectivity matrix computation

To reconstruct the functional connectome, the brain needs to be divided into a set of nodes. Network edges connecting pairs of nodes are then defined as a measure of relationship between their respective time courses. In this work, we used the Brainnetome atlas (Fan et al., 2016), a cross-validated connectivity-based parcellation scheme that divides the cerebrum into 210 cortical and 36 subcortical gray matter regions, taken as network nodes. To reflect the main dimension of signal variation across each region (Friston et al., 2006), the first eigenvariates of the time series of all voxels included in each region mask were extracted using *fslmeants* and taken as the time course of the corresponding nodes.

Individual 246 × 246 connectivity matrices were then reconstructed by calculating the standardized regression coefficient between the time courses of nodes *i* and *j*, while also entering the ICA-AROMA artifact time courses, six motion parameters, and white matter and cerebrospinal fluid mean time courses as regressors. This produces fully connected and undirected brain networks with 30, 135 unique edges with values ranging between −1 and 1.

Moreover, a group temporal-concatenation spatial ICA was performed using FSL’s MELODIC (https://fsl.fmrib.ox.ac.uk/fsl/fslwiki/MELODIC) with a pre-determined dimensionality of 15 independent components. The spatial and power spectral characteristics of the resulting components were inspected to identify 10 well-established networks (Griffanti et al., 2017): two primary visual networks (PVN and PVN2), primary-secondary visual network (PSVN), default mode network (DMN), dorsal attentional network (DAN), anterior default mode network (aDMN), right and left frontoparietal networks (FPN), sensory-motor network (SM) and the salience network (SN) (see Supplementary Figure 1). Two additional networks were identified as the medial temporal network (MTN) (including the hippocampus and the amygdala) and the striatal-thalamic network (STN).

These data-driven spatial maps were used to assign the Brainnetome’s nodes to specific brain networks for further regional graph-based measurements calculation, using in-house MATLAB scripts. Specifically, the Z-maps generated by MELODIC corresponding to the 12 networks of interest were thresholded at Z ≥ 2. Subsequently, each Brainnetome node was considered to belong to a network if over 60% of its voxels overlapped with this network’s thresholded map. When this overlap occurred for more than one network, the sum of the Z-values across all overlapping voxels was considered, and the node was assigned to the network with the highest Z-value sum. In total, 153 of the 246 Brainnetome nodes were ascribed to one of the 12 networks of interest, as shown in Supplementary Table 1.

### 2.5. Global and regional graph measurements estimation

Global network parameters were computed using the Brain Connectivity Toolbox (Rubinov & Sporns, 2010) and in-house MATLAB scripts. After setting negative edge weights to zero, normalized weighted global measures were computed by averaging the ratio between the actual measure and those obtained through randomly rewiring the connectivity matrix 500 times using Maslov-Sneppen’s degree-preserving algorithm (L_*norm*_ = L / L_*rand*_ and C_*norm*_ = C / C_*rand*_). These normalized metrics were the characteristic path length, defined as the average minimum number of edges that need to be traversed between each pair of network nodes, and the clustering coefficient, here computed as the average of existing connections between nodes divided by the number of possible connections. Additionally, we calculated the SW index as the ratio between these abovementioned measures (SW = C_*norm*_ / L_*norm*_). Finally, we computed the mean network degree (i.e., average number of positive connections linked to a node, across all brain nodes) and the regional degree of nodes ascribed to each ICA-based network and the rest of the brain.

### 2.6. Statistical analysis

Sociodemographic variables as well as the mean interframe translational and rotational movements of the head were examined across groups. Non-parametrical tests were applied when data assumptions were violated. All comparisons, including post-hoc pairwise contrasts, of both global and regional graph-based measures were conducted with permutation-based tests (10,000 permutations) (*permuco* package; Renaud and Frossard, 2019).

One-factor univariate general linear models (GLMs) were conducted in adults and seniors. Post-hoc pairwise t-tests were performed for contrasts that emerged as statistically significant (*RVAideMemoire* package; Herve, 2009). Cumulative odds in post-hoc comparisons for false-positive results were prevented with Bonferroni-adjustments. Statistical significance was set at Bonferroni-adjusted two-tailed p-value < 0.05. Likewise, effect sizes (i.e., Cohen’s d [d]) and confidence intervals (CI) at 95% were calculated for each significant contrast with bootstrap procedures (10,000 simulations) (bootES package; Gerlanc and Kirby, 2015). All statistical analyses were performed with R version 3.5.0 (R Core Team, 2018).

## 3. RESULTS

### 3.1. Sociodemographic

As expected, groups differed in age (F_(2,90)_ = 329, p < 0.001) and years of education (F_(2,90)_ = 10.85 p < 0.001). Post-hoc tests revealed no significant differences between adults with and without obesity in age (p-adjusted = 0.428) or education (p-adjusted = 0.371). All groups were equally distributed for sex (x^2^ = 1.82, p = 0.402). Please see Table 1 for a summary of demographic variables.

**Table 1.** Sociodemographics across groups (mean and standard deviation, or %)

|  | N | Females | Age | Range | Education | Range |
| --- | --- | --- | --- | --- | --- | --- |
| **OB adult** | 30 | 20 (66.7%) | 32.8 (5.7) | 21 – 39 | 13.7 (2.9) | 10 – 20 |
| **HW adult** | 33 | 18 (54.5%) | 30.9 (6.2) | 21 – 40 | 14.8 (2.5) | 10 – 20 |
| **Senior** | 30 | 15 (50%) | 67.1 (6.7) | 56 – 85 | 11 (4.3) | 6 – 20 |

### 3.2. MRI quality-check

Rotational movements of the head did not diverge between groups (H_(4)_ = 4.86, p = 0.09). By contrast, head translational movements did prove different (H_(4)_ = 20.19, p < 0.001). Post-hoc contrasts revealed that while both elders and adults with obesity differed from the healthy-weight group, they did not diverge from each other. Consequently, head translational movements were controlled for all further graph-theoretical analyses.

### 3.3. Global results for graph-theoretical measurements

Groups showed differences in the SW index (F_(3,89)_ = 13.43, p < 0.001) with post-hoc analyses revealing that adults with (t = 5.33, p-adjusted < 0.001; d = 1.38; CI_bca95_% [0.70 ~ 2.03]) and without obesity (t = 2.97, p-adjusted = 0.011; d = 0.75; CI_bca_95% [0.18 ~ 1.31]) proved greater scores than elders. Bonferroni correction prevented the difference between individuals with obesity and controls to be statistically significant (raw p-value = 0.045).

In-depth analysis of these SW index deviations returned group differences in the clustering coefficient (F_(3,89_ = 10.02, p < 0.001), but not in the characteristic path length (F_(3,89)_ = 2.31, p = 0.105). Here, only the obesity group showed a higher clustering coefficient relative to seniors (t = 4.76, p-adjusted < 0.001; d = 1.23, CI_bca_95% [0.64 ~ 1.88]). The contrast between controls and seniors did not survive Bonferroni adjustment (raw p-value = 0.037). The comparison including adults with and without obesity did not pass multiple comparison correction either (raw p-value = 0.023).

Moreover, the mean network degree assessments uncovered group differences (F_(3,89)_ = 4.99, p = 0.008). Here, participants with obesity demonstrated a lower average number of effective connections as compared to seniors (t = −2.96, p-adjusted = 0.013; d = −0.77; CI_bca_95% [−1.26 ~ −0.26]) and healthy-controls (t = −2.89, p-adjusted = 0.020; d = −0.72; CI_bca_95% [−1.19 ~ −0.27]). No differences between controls and seniors were found.

### 3.4. Regional nodal degree and strength results

Groups diverged for the average number of inward and outward connections across several functional networks, namely the DMN (F_(3,94)_ = 6.82, p-value = 0.002), the DAN (F_(3,94)_ = 4.63, p-value = 0.012), the SM (F_(3,94)_ = 5.17, p-value = 0.009), the left FPN (F_(3,94)_ = 3.63, p-value = 0.031), the SN (F_(3,94)_ = 3.48, p-value = 0.030) and the PSVN (F_(3,94)_ = 3.55, p-value = 0.034).

Post-hoc comparisons exposed that adults with obesity displayed lesser regional degree within the DMN and the rest of the brain when compared to seniors (t = −3.25, p-adjusted = 0.004; d = −0.77; CI_bca_95% [−1.23 ~ −0.22]) and healthy-weight adults (t = −3.34, p-adjusted = 0.002; d = −1.01; CI_bca_95% [−1.46 ~ −0.50]). This very same pattern emerged in this group for the DAN, again, relative to elders (t = −2.80, p-adjusted = 0.023; d = −0.60; CI_bca_95% [−1.13 ~ −0.07]) and controls (t = −2.62, p-adjusted = 0.027; d = −0.85; CI_bca_95% [−1.37 ~ −0.30]). Separately, adults with obesity showed lower regional degree when matched with controls in the SM (t = 2.83, p-adjusted = 0.010; d = −0.95; CI_bca_95% [−1.46 ~ −0.39]) and the PSVN (t = 2.40, p-adjusted = 0.023; d = −0.78; CI_bca_95% [−1.28 ~ −0.31]). By the same token, this group proved fewer effective connections than seniors in the left FPN (t = 2.78, p-adjusted = 0.018; d = −0.69; CI_bca_95% [−1.13 ~ −0.18]) and the salience network (t = 2.84, p-adjusted = 0.024; d = −0.59; CI_bca_95% [−1.15 ~ −0.03]). The multiple comparison correction was too restrictive for the contrasts between adults with obesity and seniors for the PSVN (raw p-value = 0.041), as well as for those among controls and seniors in the left FPN (raw p-value = 0.031) and the SM (raw p-value = 0.019). For a summary of global and regional comparison between group please see Figure 1 below.

**Figure 1.**
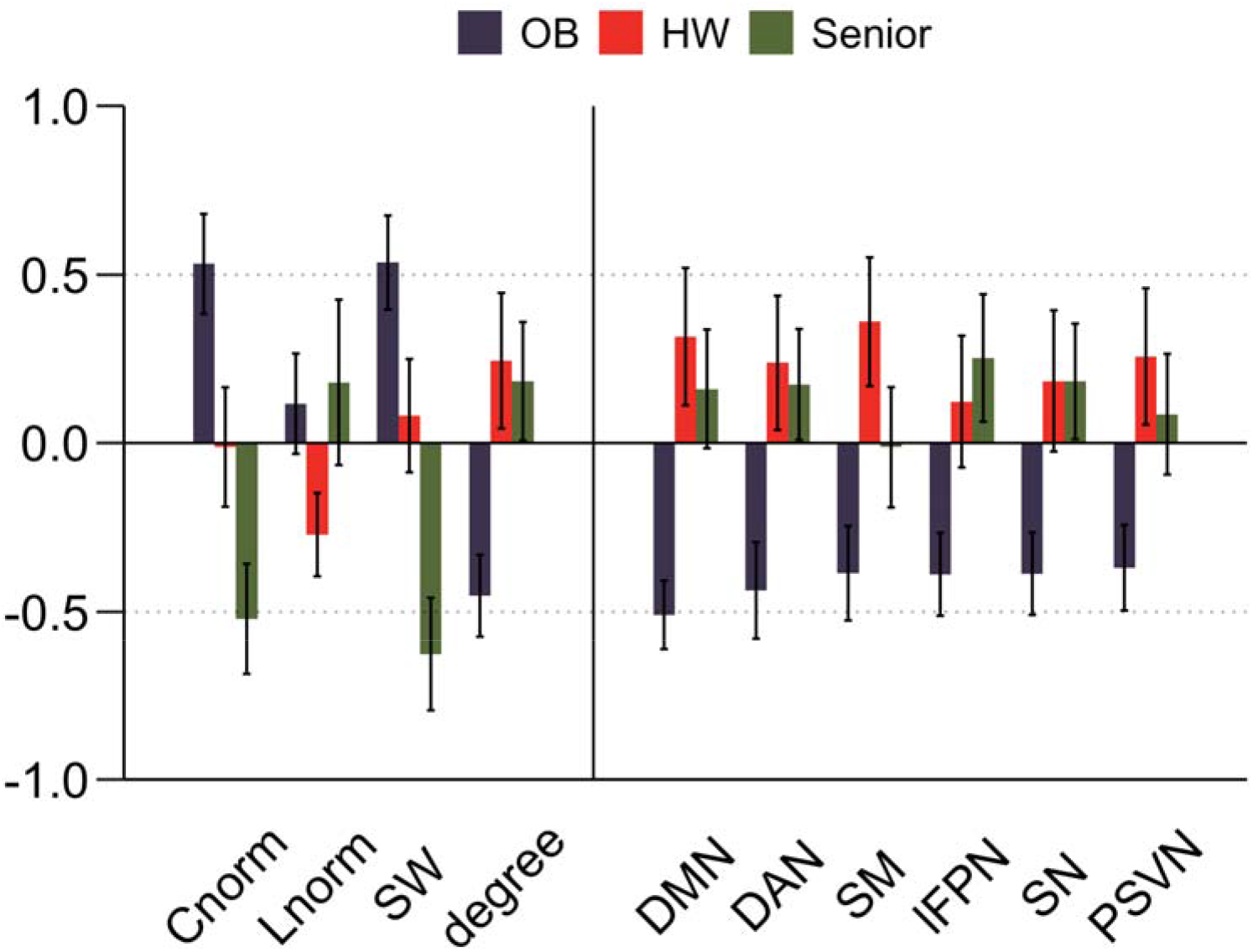
Group comparison in global and regional network metrics. OB = Adult with obesity, HW = Healthy-weight adult, Cnorm = Normalized clustering coefficient, Lnorm = Normalized characteristic path length, SW = Small-world index, DMN = Default Mode Network, DAN = Dorsal-Attentional Network, SM = Sensory-Motor Network, lFPN = Left Frontoparietal Network, SN = Salience Network, PSVN = Primary-Secondary Visual Network.

## 4. DISCUSSION

Obesity is a preventable health problem associated with the development of long-lasting cardiometabolic disorders (Wade et al., 2018) and late-onset dementia (Bischof & Park, 2015; Deckers, van Boxtel, Verhey, & Köhler, 2017). The current work aimed to shed a broader light on whether an obesity status could be related to a premature aging of the brain as formerly suggested in structural studies (Ronan et al., 2016; Tzanetakou et al., 2012). We expected to uncover a pattern mirroring the typical network rendering seen in the elderly. Contrary to what was initially postulated, adults with obesity differed from elders. Subjects with obesity demonstrated greater SM indexes that later proved being led by increases in the clustering coefficient. Moreover, individuals with obesity exhibited reductions in global and regional degree as relative to seniors and controls. The aging process is portrayed by alterations within the SM properties, mostly prompted by drops in segregation (Onoda and Yamaguchi, 2013; Brier et al., 2015; Geerlings et al., 2014; Song et al., 2014). Here, seniors partially followed this pattern as they showed reductions in their clustering coefficient as compared to adults with obesity and, to some extent, normal weight individuals.

Regarding the main objective of the study, strikingly, participants with obesity exhibit differences in the SM index when matched against seniors. Complementary analysis revealed that this was led by increases in the clustering coefficient of the obesity group. This pattern of enhanced cliquishness not only disagree with our hypothesis but also with previous obesity work (Baek et al., 2017; Chao et al., 2018) where subjects with obesity demonstrated decreases in network segregation and a shift in the network to a random-like one. These studies, however, included older subjects than the current work as well as individuals diagnosed with an eating disorder or seeking for treatment (i.e., bariatric surgery) while presenting metabolic comorbidities. Apart from these general aspects, the usage of different atlases or additional adjustments for head motion could have led to different results. A recent work have raised how weight loss significantly decreased head motion artifacts during an MRI acquisition (Beyer et al., 2020), therefore suggesting that such condition is linked to undesired movements inside the scanner. Graph-based parameters are also susceptible of the scanning protocol and the image preprocessing pipeline. Hence, additional research is needed to clarify whether an obesity status is linked to increases in network segregation.

The increases in network segregation seen in the obesity group were, however, not balanced with improvements in integration. Now, an overly clustered brain could be indicative of an ongoing drift towards a regular-like type of network. As early mentioned, a typical SM network is known for considering conflicting demands of segregation and integration. On the contrary, regular networks are notorious for being highly segregated but poorly integrated. This network architecture is often seen in some psychiatric and neurological diseases (Liao, Vasilakos, & He, 2017) as well as in very impulsive persons (Davis et al., 2013). Ultimately, this rendering could represent a flaw for superior cognitive functions (Sporns, 2013). An imbalance between the number of within and between-modular edges could translate into increases in the processing times or in a poorer performance when challenged with highly demanding tasks (Cohen & D’Esposito, 2016). These two circumstances are common findings in obesity (Deckers et al., 2017). While the average shortest path length was similar among elders and adults with obesity, this measure was, qualitatively at least, greater in the obesity group as to healthy controls. Nevertheless, this contrast did not emerge as statistically significant.

Relative to the global mean degree results, adults with obesity revealed a lower number of effective connections when rivalled to both controls and seniors. These results are partially in line with the study conducted by Geha et al. (2017). In this work, the authors presented a measure of global brain connectivity, defined as the number of connections each node has with others, which is reasonably similar to a measure of network degree. As mentioned above, one of the consequences of aging on the brain is the progressive trimming of nodes and edges (Geerligs et al., 2015; Sala-Llonch et al., 2015, 2014). Taken together with the increases in segregation, this result could suggest that, as compared to short-range links, adults with obesity may have fewer long-range connections. By contrast, degree decreases in seniors could obey instead to an excessive trimming of within-modular edges.

What is more, when examined regionally, we found group differences in sensory-driven (i.e., SM and PSVN), task-negative (i.e., DMN and SAL) and goal-oriented (i.e., lFPN and DAN) networks. On average, adults with obesity presented fewer connections within and between these networks and the rest of the brain. The functional properties of sensory-processing circuits were found to linearly decrease with age (Zonneveld et al., 2019). Other work have also reported that subjects with obesity feature alterations within tactile and visual-processing circuits (Doucet et al., 2017; García-García et al., 2013; Geha et al., 2017), and also saliency-dependent networks (García-García et al., 2013). Moreover, the DMN, whose alterations are considered as a biomarker for unsuccessful aging (Buckner, Andrews-Hanna, & Schacter, 2008; Palmqvist et al., 2017), is extremely vulnerable to the negative effects of obesity comorbidities such as diabetes (Wang, Ji, Lu, & Zhang, 2016), hypertension (Haight et al., 2015), and dyslipidemia (Spielberg et al., 2017). To date, few studies have described alterations in the functional connectivity of the DMN relative to obesity (Beyer et al., 2017; Doucet et al., 2017; Figley, Asem, Levenbaum, & Courtney, 2016; Tregellas et al., 2011). Task-positive circuits, such as the DAN or the FPN, also experience changes while aging (Geerligs et al., 2015; Sala-Llonch et al., 2015; Zhang et al., 2014) and under an obesity condition (Geha et al., 2017). Overall, this evidence stands in line with the profile depicted in the previous paragraph. Our findings indicate that the ratio between short and long-range paths is unbalanced, and so functional networks might not be trading information in the most efficient fashion as they apparently have fewer shortcuts between them. As suggested in Doucet et al. (2017), a decrease in the crosstalk between sensory-driven, internally guided and self-regulatory networks could negatively impact feeding. This could either occur through an enhanced sensitivity to food sensory properties, increases of craving and thoughts about highly palatable meals or failed attempts to refrain from overeating.

Interestingly, healthy-weight adults also differed from seniors in very few parameters. Apart from the deviations in the SM index, we failed to replicate the negative association between age and network segregation described in the literature (Geerligs et al., 2015; Song et al., 2014). Still, it is noteworthy to point that the Bonferroni-adjustment to prevent false positive results could have been particularly conservative in this case (raw p-value 0.037). Likewise, no differences in integration were reported. Still, the number of edges to cross from one module to another was qualitatively longer in elders. That said, we cannot rule out the possibility that the current study was underpowered to catch subtle group differences as the ones we found failed to pass additional thresholds. The same statistical limitation applies to the regional degree differences that most networks showed when comparing elders and controls. Moreover, it is important to remark that none of the participants in the senior group was obese, a condition that alone (Beyer et al., 2017; Doucet et al., 2017), or in conjunction with its medical complications (Haight et al., 2015; Spielberg et al., 2017; Wang et al., 2016), proved enough to yield differences in brain connectivity. This criterion is often overlooked in healthy-aging literature and should be considered onwards.

The current work has limitations that need to be addressed in the future. First, being this a transversal study, concluding upon causality is discouraged by all means. Longitudinal approaches would help disentangle whether the transition from adulthood to elderhood is different in the presence of an unhealthy weight. Second, our modest sample size (N = 93) might have limited our statistical power to retrieve small effects, if any. Third, to have had a group of elders with obesity would have allowed exploring the impact of such condition across all possible age ranges. By contrast, two strong points of this work were the efforts in controlling type I errors and the exclusion of participants with neurological or psychiatric comorbidities.

## 5. CONCLUSIONS

In sum, compared to non-obese seniors, volunteers with obesity exhibited increases in network segregation. Relative to healthy-weight adults and elders, this group also revealed important global and regional reductions in the amount of connections. Overall, an obesity status may represent an avoidable condition negatively impacting network topology. Although more research is needed, this overly clustered rendering is the opposite to the typical changes seen in elders. This network organization may in turn represent a shortcoming for superior cognitive functions that depends on the rapid interaction between different functional communities.

## 6. Acknowledgments

The authors would like to thank all the participants and collaborators who made this study possible. This work was supported by the Spanish Ministry of Economy, Industry and Competitiveness (PSI2013-48045) and the Spanish Ministry of Science, Innovation and Universities (PSI2017-8653). None of the authors had any biomedical or financial interests that might potentially bias their work. The data that support the findings of this study are available from the corresponding author upon reasonable request.

